# *KCNH2* encodes a nuclear-targeted polypeptide that mediates hERG1 channel gating and expression

**DOI:** 10.1101/2022.08.24.504830

**Authors:** Abhilasha Jain, Olivia Stack, Saba Ghodrati, Francisco G. Sanchez-Conde, Chiamaka Ukachukwu, Shreya Salwi, Eric N. Jimenez-Vazquez, David K. Jones

## Abstract

*KCNH2* encodes hERG1, the voltage-gated potassium channel that conducts the rapid delayed rectifier potassium current (IKr) in human cardiac tissue. hERG1 is one of the first channels expressed during early cardiac development, and its dysfunction is associated with intrauterine fetal death, sudden infant death syndrome, cardiac arrhythmia, and sudden cardiac death. Here, we identified a novel hERG1 polypeptide (hERG1NP) that is targeted to the nuclei of immature cardiac cells, including hiPSC-CMs and neonatal rat cardiomyocytes. The nuclear hERG1NP immunofluorescent signal is diminished in matured hiPSC-CMs and absent from adult rat cardiomyocytes. Antibodies targeting distinct hERG1 channel epitopes demonstrated that the hERG1NP signal maps to the hERG1 distal C-terminal domain. *KCNH2* deletion using CRISPR simultaneously abolished IKr and the hERG1NP signal in hiPSC-CMs. We then identified a putative nuclear localization sequence (NLS) within the distal hERG1 C-terminus, 883-RQRKRKLSFR-892. Interestingly, the distal C-terminal domain was targeted almost exclusively to the nuclei when overexpressed HEK293 cells. Conversely, deleting the NLS from the distal peptide abolished nuclear targeting. Similarly, blocking α or β1 karyopherin activity diminished nuclear targeting. Finally, overexpressing the putative hERG1NP peptide in the nuclei of HEK cells significantly reduced hERG1a current density, compared to cells expressing the NLS-deficient hERG1NP or GFP. These data identify a developmentally regulated polypeptide encoded by *KCNH2*, hERG1NP, whose presence in the nucleus indirectly modulates hERG1 current magnitude and kinetics.

## INTRODUCTION

The KCNH2 gene encodes hERG1, the voltage-gated potassium channel that conducts the critical cardiac repolarizing current, IKr (1, 2). KCNH2 variants that reduce IKr, or off-target hERG1 channel block cause the cardiac disorder long QT syndrome (LQTS) and increase the likelihood for cardiac arrhythmia and sudden cardiac death (2, 3). KCNH2 variants have also been linked with a number of causes of death in the young that may or may not be directly linked to cardiac dysfunction, including intrauterine fetal death (4), sudden infant death syndrome (SIDS) (5, 6), and sudden unexplained death in epilepsy (SUDEP) (7–9).

The KCNH2 gene encodes multiple hERG1 splice variants in cardiac tissue (10–12). Conducting hERG1 channels comprise at least two subunits encoded by alternate KCNH2 transcripts, hERG1a and hERG1b, that are identical apart from their N-terminal domains (10, 11). hERG1a contains an N-terminal Per-Arnt-Sim (PAS) domain that regulates gating through dynamic interactions with the S4-S5 linker and the cyclic nucleotide binding homology domain (CNBHD) of the proximal C-terminus (13–17). hERG1b has a much shorter and unique N-terminus that lacks a functional PAS domain (10, 11). When expressed in HEK293 cells, the absence of a PAS domain in hERG1b accelerates the time course of activation, deactivation, and inactivation recovery of heteromeric hERG 1a/1b channels by two-fold, compared to homomeric hERG1a channels (18). In human cardiomyocytes, silencing hERG1b by overexpressing a polypeptide that mimics the hERG1a PAS domain slows native IKr gating kinetics and reduces IKr magnitude, prolonging the action potential duration (19). Conversely, disabling the hERG1a PAS domain using PAS-targeting antibodies accelerates IKr gating, increases IKr magnitude, and hastens cardiac repolarization (20). KCNH2 also encodes a non-conducting C-terminal splice variant that reduces total hERG1 current density, hERG1USO (12, 21, 22). hERG1USO can combine with either N-terminal variant (hERG1a vs hERG1aUSO, & hERG1b vs hERG1bUSO) (12, 21, 22). The relative abundance of hERG1 subunits shifts during fetal development (4) and heart failure (23), suggesting that hERG1 subunit abundance is dynamic.

The hERG1 C-terminus is a critical component of hERG1 channel function. The proximal C-terminus (670-872, hERG1a numbering) is highly structured and contains a C-linker and a cyclic nucleotide binding homology domain (CNBHD) that play integral roles in channel gating (13, 15, 17, 24–29). Mutations throughout the C-linker and CNBHD trigger dramatic changes in hERG1 gating and surface expression (24, 30). In contrast, the distal C-terminal domain (863-1159, hERG1a numbering) is disordered (31, 32) and its contribution to hERG1 function is less clear. In fact, several SIDS-associated hERG1 variants in the distal C-terminus have a limited impact on hERG1 gating or expression (33). Thus, the causal link between many KCNH2 variants and sudden death in the young remains unclear and the regulatory mechanisms of hERG1 in immature cardiomyocytes is underexplored.

To identify novel regulators of hERG1 expression in developing cardiomyocytes we leveraged the immature nature of human stem cell-derived cardiomyocytes (hiPSC-CMs). hiPSC-CMs display maturation comparable to cardiomyocytes isolated from human embryonic and/or fetal cardiac tissue (34). And with the increasing accessibility of CRISPR technology, hiPSC-CMs are a uniquely powerful tool to explore regulatory mechanisms in immature human cardiomyocytes. Here we identify a non-conducting KCNH2-encoded polypeptide, hERG1NP, that is targeted to the nuclei of immature cardiac cells, including hiPSC-CMS and neonatal rat cardiomyocytes, but not adult rat cardiomyocytes. hERG1NP expression within the nucleus reduced hERG1 current density by depolarizing the voltage-dependence of activation and stabilizing inactivation. These data identify hERG1NP as a developmentally regulated, KCNH2-encoded polypeptide that regulates hERG1 function via the cell’s nucleus.

## RESULTS

### A hERG1 Immunofluorescence Signal in Cardiac Nuclei

To identify regulatory mechanisms of hERG1 function, we cultured human pluripotent stem cell-derived cardiomyocytes (hiPSC-CMs) on Matrigel-coated glass coverslips. We used phalloidin labeling actin to identify cardiomyocytes from other cell types, as previously described (19, 35). We immunolabeled for hERG1 using antibodies targeting two distinct epitopes: (1) the extracellular p-loop and (2) the distal C-terminal domain (Fig. 1a). We then measured immunofluorescence using confocal microscopy to track the distribution of the two antibodies. Surprisingly, the immunolabeling pattern of the two antibodies did not overlap. The distribution of the p-loop-targeting antibody was enriched in the cardiac ER, cytoplasmic space, and surface membrane, consistent with previous reports of hERG1 distribution in cardiac cells (36–39). In contrast, the antibody targeting the hERG C-terminal domain was enriched within the nuclei (Fig. 1b). To quantify the relative distribution of nuclear vs non-nuclear hERG1 fluorescence, we divided the mean hERG1 fluorescence intensity within the nucleus by the mean fluorescence intensity within the cytoplasm from each cell. The ratio of nuclear to cytoplasmic fluorescence was significantly higher for the C-terminal antibody compared to the p-loop antibody (p < 0.0001) (Fig. 1c). These data suggest that hiPSC-CMs target a portion of the hERG1 protein to their nuclei.

**Figure 1.**
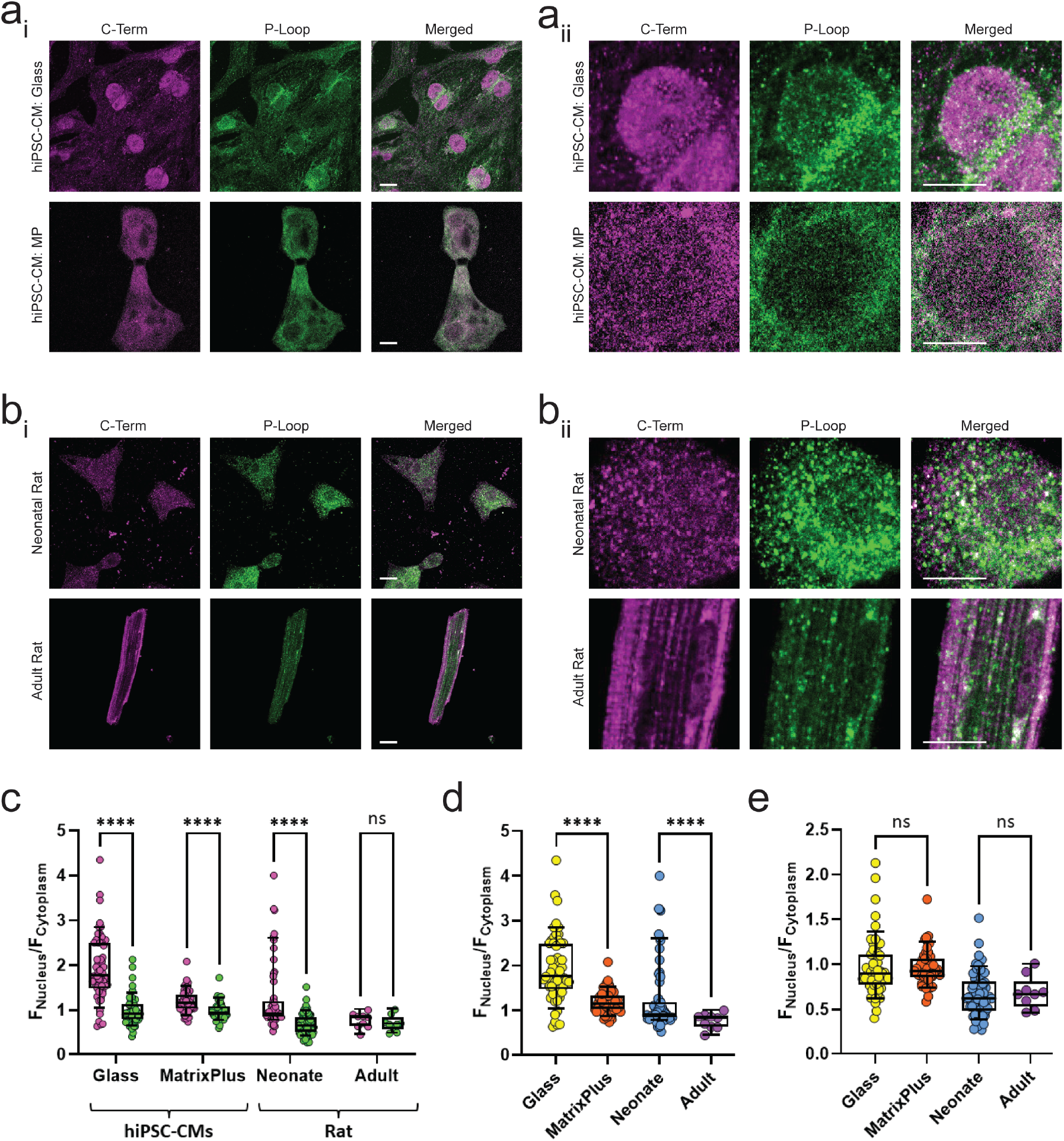
ERG1 Immunofluorescence Signal in Cardiac Nuclei. (a_i_ & a_ii_) hiPSC-CMs cultured on glass (top) or Matrixplus (bottom). Cells are dual labeled for hERG1 with antibodies targeting the carboxy terminus (C-Term) and the extracellular pore (P-Loop). (a_ii_) Expanded view of select nuclei from images in “ai”. (b_ii_ & b_ii_) Cardiomyocytes of neonatal rat (top) or adult rat (bottom) labeled for rERG1 with antibodies as in “a”. (b_ii_) Expanded view of select nuclei from images in “bi”. Scale bars indicate 10 μM. (c) Box and whisker plots depicting nuclear fluorescence intensity relative to cytoplasmic fluorescence intensity (FNucleus/FCytoplasm) for the C-Term and P-loop antibodies. The C-Term antibody was enriched in the nucleus compared to the P-Loop antibody in immature cardiomyocytes, but not adult cardiomyocytes. (d) Box and whisker plots demonstrating that the relative abundance of nuclear ERG1 fluorescence of the C-Term antibody (FNucleus/FCytoplasm) decreased with increased cardiac maturation. (e) The intracellular distribution of the P-Loop antibody was unaffected by cardiomyocyte maturation. **** indicates p < 0.0001.

### Nuclear hERG1 Distribution Corresponds with Cardiac Maturation

To determine if these results were an artifact of the in vitro differentiation process, we repeated these experiments in freshly isolated neonatal and adult rat cardiomyocytes (Fig. 1b). Antibodies targeting the rERG1 p-loop and C-terminal domain in neonatal rat cardiomyocytes displayed a similar distribution as hERG1 in hiPSC-CMs, where the C-terminal antibody displayed greater nuclear enrichment compared to the p-loop antibody. Interestingly, adult rat myocytes did not display distinct distribution of the p-loop and C-terminal rERG1 antibodies (Fig. 1c), and nuclear rERG1 staining was significantly lower in adult rat compared to neonatal rat cardiomyocytes (Fig. 1d). Importantly, the distribution of the p-loop antibody was unchanged between neonatal and adult rat cardiomyocytes (Fig. 1e). These data suggest that the abundance of nuclear ERG1 is dependent upon the maturation state to the cardiomyocyte.

Cardiomyocytes differentiated from human stem cells and cultured on Matrigel-coated coverslips display characteristics consistent with those observed in human embryonic/fetal cardiomyocytes, including poor sarcomere organization, low cTnI expression, and depolarized resting membrane potentials (19, 34, 40, 41). We evaluated if in vitro hiPSC-CM maturation could reduce the nuclear distribution of the C-terminal hERG1 signal, similarly to that observed between neonatal and adult rat cardiomyocytes. To induce hiPSC-CM maturation we cultured hiPSC-CMs on a commercial cardiac maturation matrix, MatrixPlus (42). hiPSC-CMs cultured on MatrixPlus display rod-shaped morphology, highly organized sarcomeres, elevated cTnI expression, and resting membrane potentials at or near −80 mV (42). In MatrixPlus-cultured hiPSC-CMs, the C-terminal antibody displayed greater nuclear enrichment compared to the p-loop antibody (Fig. 1c & d). However, nuclear hERG1 enrichment was significantly reduced in MatrixPlus-cultured hiPSC-CMs compared to hiPSC-CMs cultured on glass, and more closely resembled rERG1 levels measured from rat neonatal cardiomyocytes (Fig. 1d). The distribution of the p-loop antibody was unchanged between glass-cultured and MatrixPlus-cultured hiPSC-CMs (Fig. 1e). These data demonstrate that the abundance of the nuclear ERG1 immunofluorescent signal associated with the ERG1 C-terminus is dependent upon the maturation state of the cardiomyocyte.

### KCNH2 Deletion Abolishes the Nuclear hERG1 Immunosignal

To validate the accuracy of the hERG1 nuclear immunofluorescent signal, we immunolabeled for hERG1 in a line of stem cell-derived cardiomyocytes where KCNH2 expression was abolished by CRISPR (Fig. 2). Isogenic control and KCNH2-null stem cells displayed markers of pluripotency (Fig. 2a), and successfully differentiated into contracting cardiomyocytes displaying the reticulated actin staining corresponding with alignment of the cardiac sarcomeres (Fig. 2b). Isogenic control cardiomyocytes conducted IKr and displayed both cytoplasmic and nuclear hERG1 immunosignals using the C-terminal-targeting antibody. In contrast, KCNH2 double null cardiomyocytes cultured on Matrigel-coated glass lacked IKr and did not display a discernable hERG1 immunosignal in the cytoplasm or nucleus (Fig. 2b-e). These data unequivocally validate the specificity of hERG1 immunofluorescent signal in the cardiac nucleus and identify a novel KCNH2-encoded polypeptide that we termed: hERG1NP.

**Figure 2.**
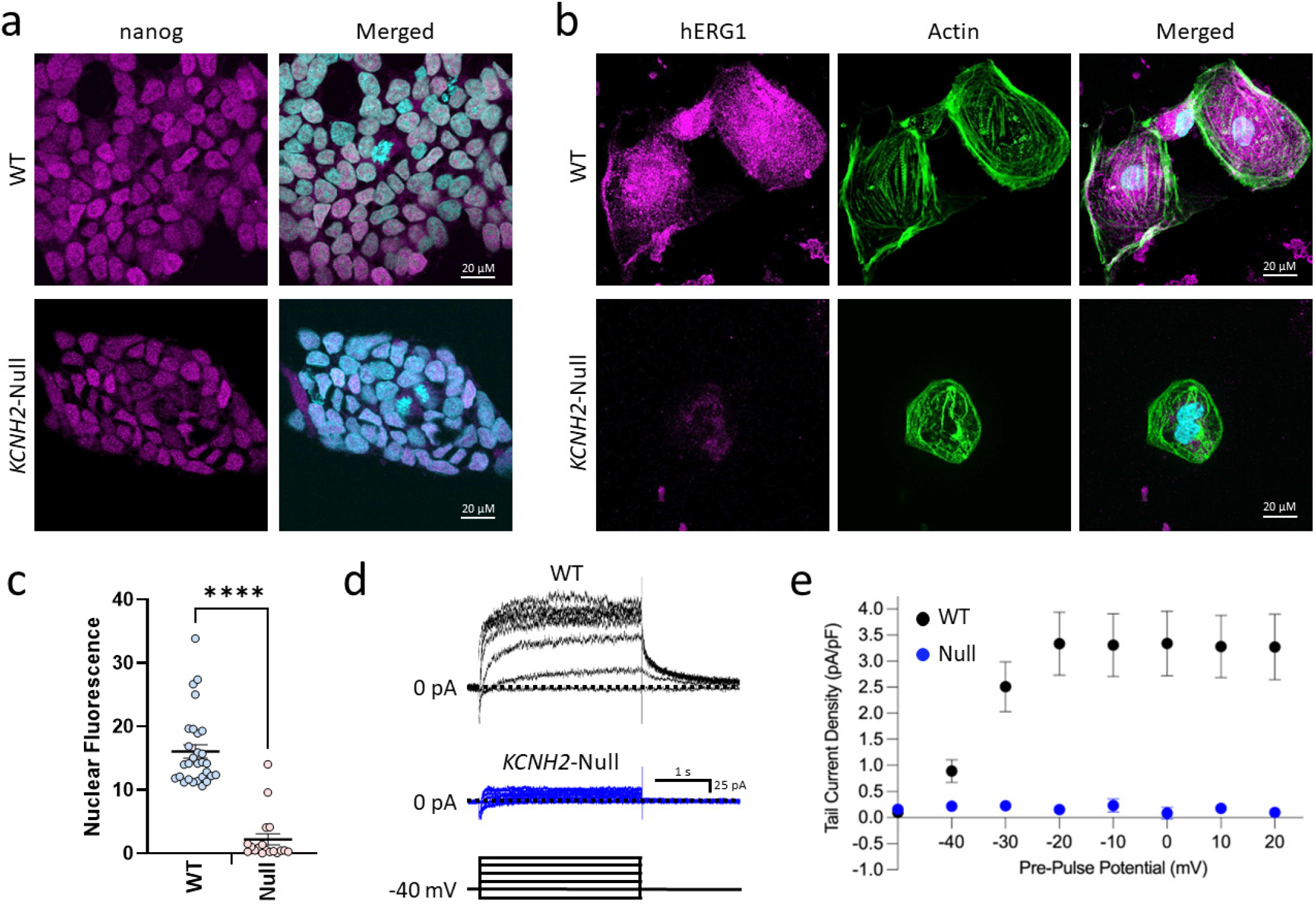
KCNH2 Deletion Abolishes the Nuclear hERG1 Signal. (a) Wildtype isogenic control (WT) and KCNH2-null hESCs stained for the pluripotency marker nanog (magenta) and DAPI (cyan). (b) Wildtype isogenic control (WT) and KCNH2-null hESC-CMs stained for hERG1 with the C-Term antibody (magenta), along with phalloidin (green) and DAPI (cyan) to mark actin and nuclei, respectively. (c) Mean nuclear fluorescence recorded from wildtype (WT, blue) or KCNH2-null (Null, red) hESC-CMs. KCNH2-null hESC-CMs do not display a nuclear hERG1 signal. (d) Sample E-4031-sensitve currents, indicative of IKr, recorded from wildtype isogenic control hES-CMS (top, black) and KCNH2-null hESC-CMs (bottom, blue). A schematic depicting the pulse protocol used is depicted at bottom. (e) Peak tail IKr density plotted as a function of pre-pulse potential for wildtype isogenic control (black) and KCNH2-null (blue) cardiomyocytes. KCNH2 null hES-CMs did not conduct a discernable IKr. n = 9 and 6 for wildtype and KCNH2-null IKr recordings, respectively. Data are represented as mean ± standard error. **** indicates p < 0.0001.

### The α/β1 Karyopherin Complex Regulates hERG1NP Nuclear Transport

The complex between the α and β1 karyopherins mediate the classical nuclear transport pathway (43). To assess if hERG1NP trafficking is dependent upon this pathway we inhibited α and β1 karyopherin activity using ivermectin and importazole, respectively (44, 45). We incubated hiPSC-CMs in 40 μM importazole for 8 hrs and then measured the relative nuclear and cytoplasmic distribution of hERG1 using the C-terminal antibody as described above (Fig. 2). 40 μM importazole significantly reduced the relative intensity of the nuclear hERG1 signal compared to vehicle controls (Fig. 3b), indicating that nuclear transport of the hERG1NP is karyopherin-β1 dependent. To test if hERG1NP trafficking is also dependent upon the α karyopherin, we blocked incubated cells with 2.5 μM ivermectin for 8 hrs. Like importazole, ivermectin significantly reduced the relative intensity of the nuclear hERG1 signal, compared to vehicle controls (Fig. 3b). These data indicate that the classical nuclear transport pathway mediates hERG1NP localization in the nucleus.

**Figure 3.**
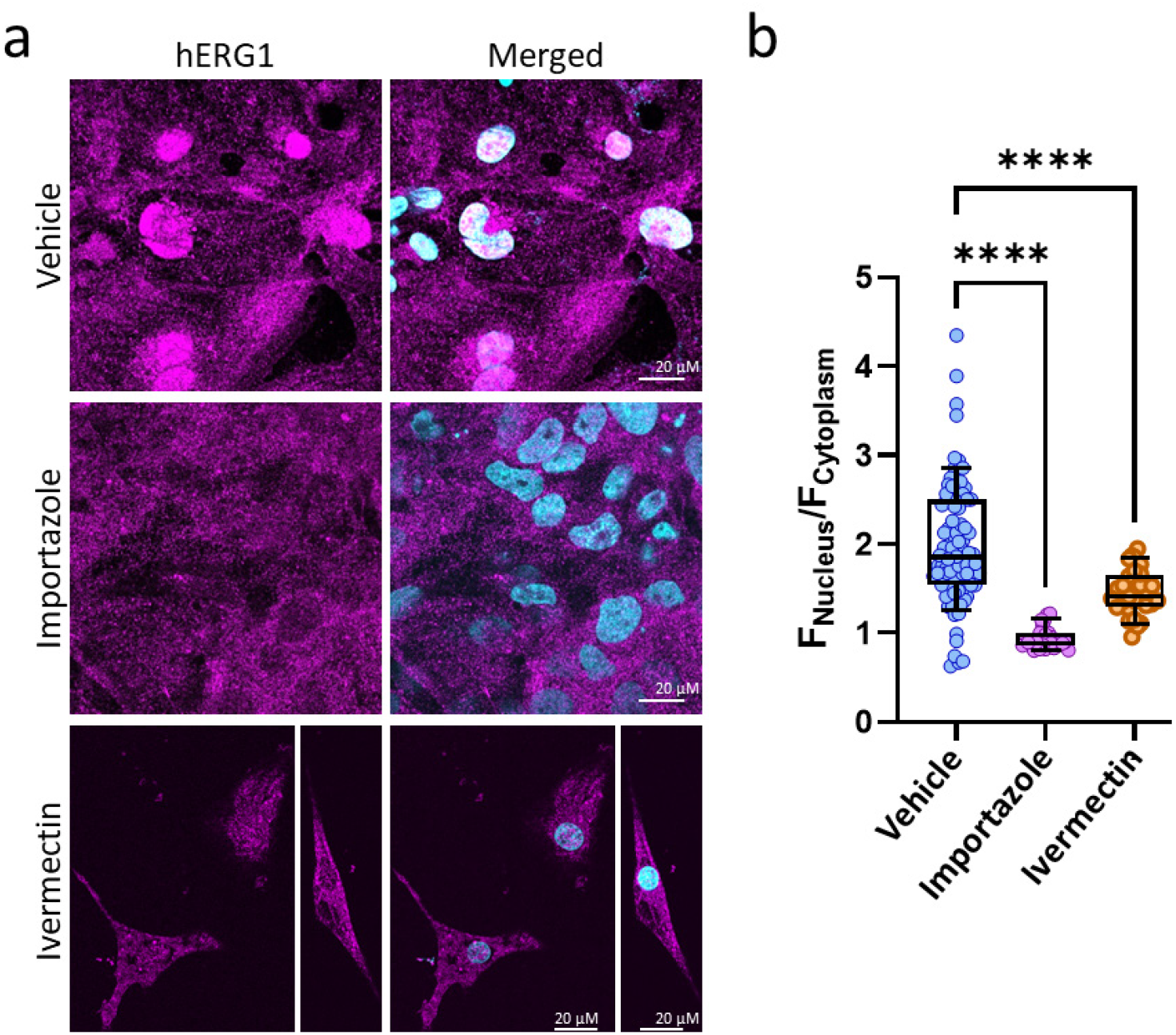
The α/β1 Karyopherin Complex Regulates hERG1NP Nuclear Transport in hiPSC-CMs. (a) hiPSC-CMs following an 8 hr incubation in either vehicle (top), 40 μM importazole (middle), or 2.5 μM ivermectin (bottom), and stained for hERG1 with the C-Term antibody (magenta) and DAPI (cyan). (b) Box and whisker plots depicting nuclear fluorescence intensity relative to cytoplasmic fluorescence intensity (FNucleus/ FCytoplasm) for the C-Term antibody. Importazole and ivermectin significantly reduced the nuclear enrichment of the C-Term antibody. **** indicates p < 0.0001

### A Nuclear Localization Sequence in the Distal C-terminal Domain of hERG1

Karyopherins recognize and bind to their cargo through monopartite or bipartite nuclear localization sequences (NLS) (46). To determine if hERG1 contains an NLS that may mediate nuclear targeting, we screened the hERG1 amino acid sequence using the open-source software cNLS Mapper. cNLS Mapper identified a single putative monopartite NLS in the distal hERG1 C-terminal domain (883RQRKRKLSFR892, Fig. 4a). This NLS is completely absent from the hERG orthologue, hERG2. The sequence is partially conserved in the hERG orthologue hERG3, however cNLS Mapper did not identify the hERG3 sequence as a potential NLS (Fig. 4b).

**Figure 4.**
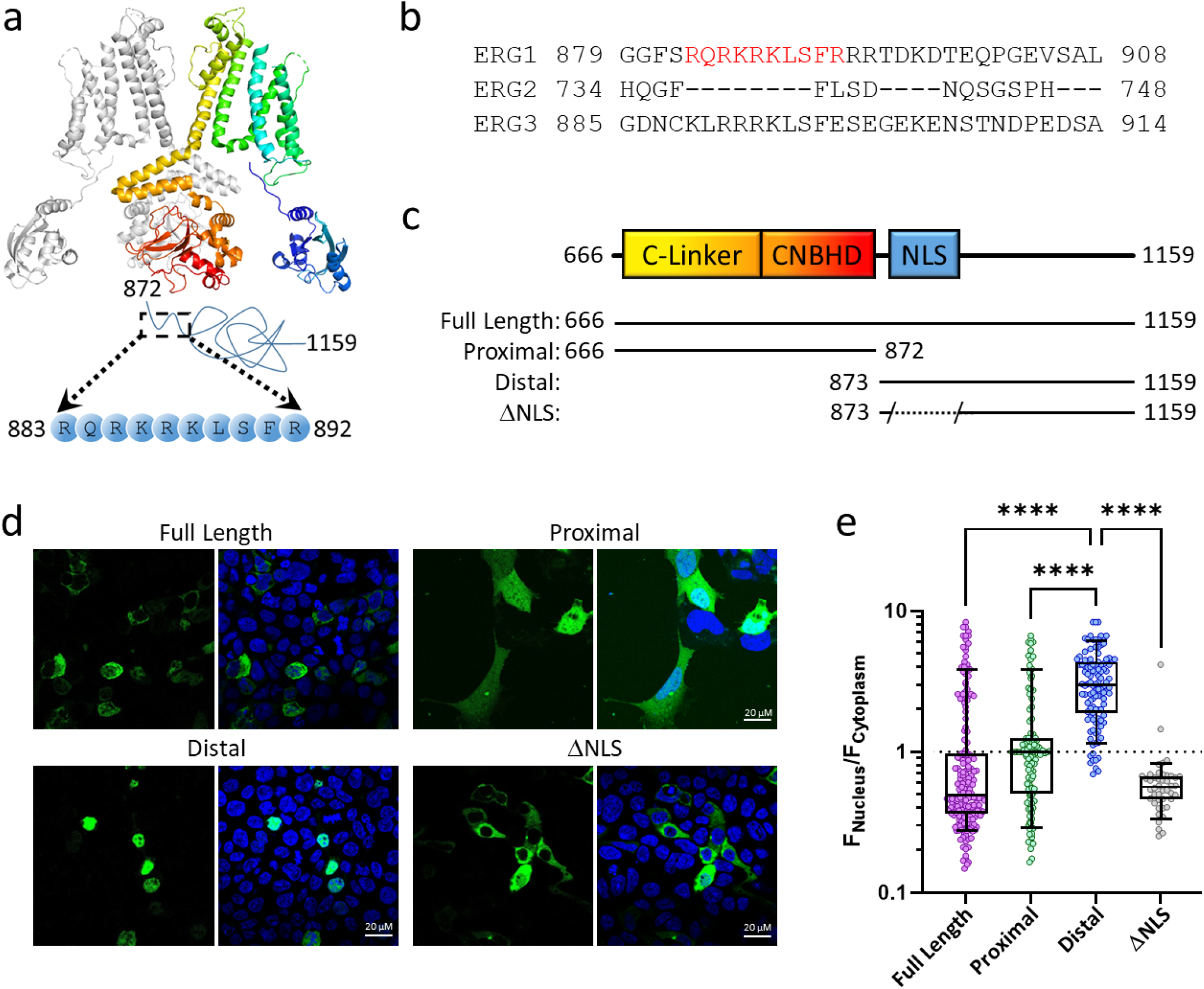
A Nuclear Localization Sequence in the Distal C-terminal Domain of hERG1. (a) Cryo-EM schematic depicting two opposing hERG1a subunits. The distal c-terminal domain with its NLS (883-892) is depicted with a blue line. (b) Amino acid sequence alignment of the ERG1 NLS with ERG2 and ERG3. (c) Schematic depicting the C-terminal coding regions of the full length (666-1159), proximal (666-872), distal (873-1159), and ΔNLS constructs (873-882/893-1159). (d) HEK293 cells expressing the full length, proximal, distal, and ΔNLS constructs. (e) Box and whisker plots depicting the intracellular distribution of HEK293 cells transfected with the full length (magenta), proximal (green), distal (blue), and ΔNLS (grey) constructs. **** indicates p < 0.0001

We then used three citrine-fused polypeptides to assess the functionality of the putative hERG1 NLS: 666-1159, 666-872, and 873-1159 (Fig. 4c). 666-1159 (full length) represents the entire hERG1 C-terminal cytoplasmic domain. 666-872 (proximal) forms the hERG1 C-linker and CNBH domains, and the 873-1159 (distal) forms the disordered distal C-terminal domain of hERG1. We expressed each fusion peptide individually in HEK293 cells and tracked the distribution of the peptides using confocal microscopy. Cells transfected with the full-length peptide (666-1159) displayed a bimodal distribution. 81% of cells displayed cytoplasmic enrichment, however a subset of cells (19%) displayed a strong nuclear enrichment (Fig. 4d, top left). The proximal peptide (666-872) lacks the NLS sequence and displayed equivalent fluorescence between the cytoplasm and nuclei (Fig. 4d, top right). In contrast, the proximal peptide (873-1159), which contains the putative NLS, was limited almost exclusively to the cells’ nuclei ((Fig. 4d, bottom left). Additionally, deleting the NLS coding region from the distal peptide construct abolished nuclear targeting (ΔNLS, Fig. 4c & d, bottom right), demonstrating that the 883RQRKRKLSFR892 sequence drives nuclear transport of the distal peptide. These results show that the hERG1 sequence 883RQRKRKLSFR892 is a functional NLS.

Finally, to determine if the nuclear targeting of the distal and full-length peptides in HEK293 cells followed the same mechanism as that observed in the hiPSC-CMs, we inhibited α and β1 karyopherin activity using ivermectin and importazole, respectively. Like the effects observed in hiPSC-CMs, both ivermectin and importazole diminished nuclear targeting of the distal peptide and abolished nuclear targeting of the full-length peptide (Fig. 5). These data demonstrate that the hERG1 NLS drives nuclear transport of the distal and full-length peptides through the classical nuclear transport pathway, identical to the native hERG1NP in hiPSC-CMs.

**Figure 5.**
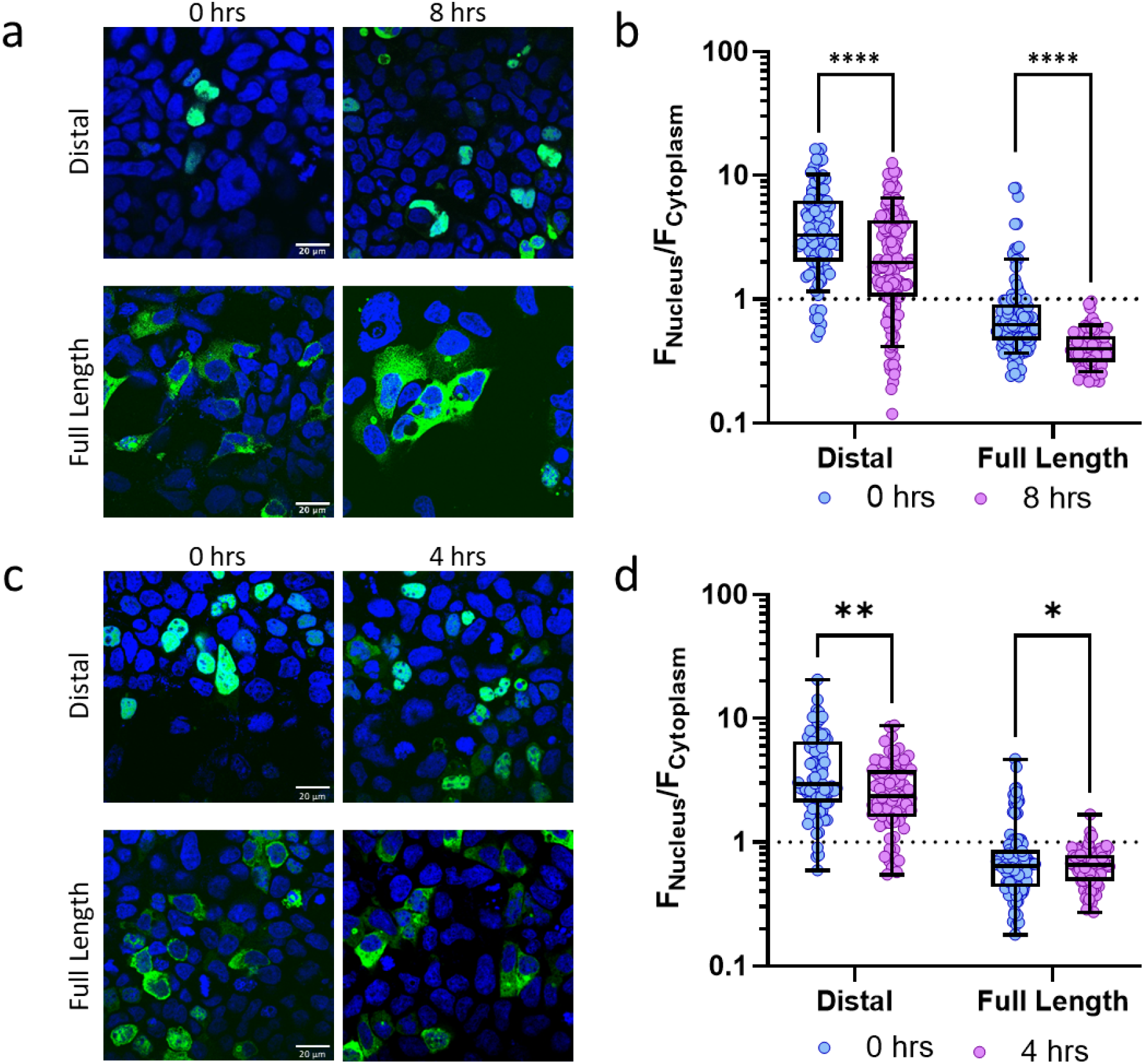
The α/β1 Karyopherin Complex regulates nuclear targeting of hERG1 C-terminal truncates. (a) HEK293 cells expressing the distal (top) and full-length (bottom) peptides (green) at 0 and 8 hrs, with 40 μM importazole. (d) Box and whisker plots depicting intracellular distribution of the full-length and distal peptides before and after 8 hr incubation with 40 μM importazole. (c). HEK293 cells expressing the distal (top) and full-length (bottom) peptides at 0 and 4 hrs, with 2.5 μM ivermectin. (d) Box and whisker plots depicting intracellular distribution of the full-length and distal peptides before and after 8 hr incubation with 2.5 μM ivermectin.* p < 0.05. indicates, ** indicates p < 0.01. **** indicates p < 0.0001

### The hERG1NP Inhibits I_hERG1_

We used the distal peptide as a tool to explore the functional impact of the hERG1NP on the electrophysiological properties of hERG1 currents. We transfected HEK293 cells stable expressing hERG1a with the distal peptide (Fig. 6). The hERG1NP reduced tail current density by 55%, compared to GFP transfected controls (Fig. 6a,b & Table 1). The hERG1NP reduced steady-state currents even further, by 78% compared to GFP controls (Fig. 6a,c & Table 1). The equilibrium between the activated and inactivated states mediates the magnitude of hERG steady-state current. We therefore hypothesized that the hERG1NP promotes hERG1 inactivation, thereby exacerbating steady-state current reduction compared to tail current reduction. To test this, we normalized steady-state currents to the maximum tail current recorded from the same cell to calculate the relative steady-state current – which is a proxy for the stability of the inactivated state. The hERG1NP reduced the relative steady-state current magnitude from 1.00 ± 0.17 AU in the presence of GFP to 0.45 ± 0.06 AU in the presence of the hERG1NP (Table 1, Fig. 6d). These data demonstrate that the hERG1NP reduces hERG1 current by reducing channels at the membrane and by promoting inactivation.

**Figure 6.**
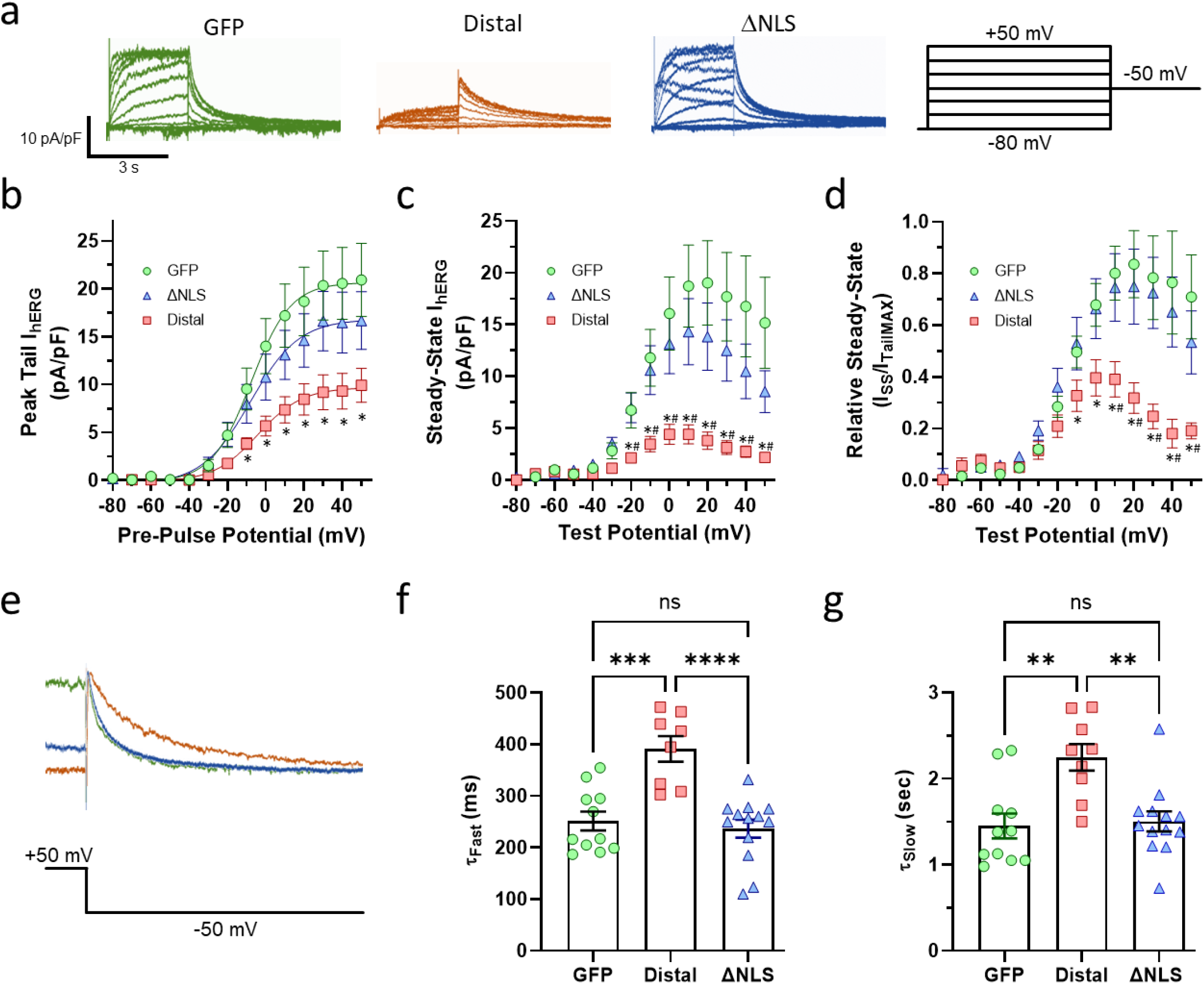
The hERG1NP Inhibits I_hERG1_. (a) Sample current traces recorded from HEK293 cells stably expressing hERG1a and transfected with vectors encoding either GFP (green), Distal (red), or ΔNLS constructs (blue). Voltage protocol used is shown at right. (b) Peak tail current density (pA/pF) plotted as a function of pre-pulse potential and fitted with a Boltzmann function, recorded from HEK293 cells stably expressing hERG1a and transfected with GFP (green circles), Distal (red squares), or ΔNLS (blue triangles). The Distal peptide significantly reduced tail current density compared to GFP and ΔNLS. (c) Steady-State current density (pA/pF) plotted as a function of test potential for GFP (green circles), Distal (red squares), or ΔNLS (blue triangles). The Distal peptide significantly reduced steady-state current density compared to GFP and ΔNLS. (d) Relative steady-state current magnitude (ISS/ITailMAX) plotted as a function of test potential for GFP (green circles), Distal (red squares), or ΔNLS (blue triangles). The Distal peptide significantly reduced relative steady-state current density compared to GFP and ΔNLS. n = 9 – 12. * indicates p < 0.05 vs GFP. # indicates p < 0.05 vs ΔNLS. (e) Sample traces depicting deactivation from GFP (green), Distal (blue), and ΔNLS (red). The distal peptide significantly slowed the time course of deactivation compared to GFP or ΔNLS. (f) Fast time constants of deactivation for GFP (green circles), Distal (red squares) and ΔNLS (blue triangles). (g) Slow time constants of deactivation for GFP, Distal and ΔNLS. ** indicates p < 0.01. *** indicates p < 0.001. **** indicates p < 0.0001.

We also measured the impact of the hERG1NP on the time course of hERG1 deactivation. hERG1NP expression significantly slowed the time course of deactivation compared to GFP-transfected controls (Fig. 6e-g). Given that the voltage dependence of activation was unaffected by the distal peptide (Table 1), these data also suggest that the distal peptide is similarly accelerating the time course of activation.

Finally, to evaluate if the effects of the distal peptide were dependent upon its targeting to inside the nucleus, we expressed the ΔNLS peptide. hERG1 currents recorded in the presence of the ΔNLS peptide were significantly larger compared to the distal peptide and similar to GFP controls (Fig. 6c-f). These data demonstrate that the effects of the hERG1NP are dependent upon its trafficking inside the nucleus. Together, these electrophysiological data demonstrate that nuclear targeting of the hERG1NP dramatically reduces hERG1 current magnitude, slows channel deactivation, and promotes hERG1 inactivation.

## DISCUSSION

Here we identified a KCNH2-encoded polypeptide, hERG1NP, which is upregulated in immature cardiomyocytes. α and β1 karyopherins, in conjunction with a nuclear localization sequence in the distal hERG1 C-terminal domain, target hERG1NP to the nucleus. Nuclear targeting of the putative hERG1NP (the distal C-terminal domain) alters hERG1a gating and dramatically reduces hERG1a current magnitude. These data demonstrate a new mechanism of regulation of hERG1.

Our study identifies two mechanisms by which that nuclear targeting of the distal peptide attenuates hERG1a currents. First, the reduced tail currents demonstrates that the distal peptide reduces hERG1a currents by attenuating hERG1a surface expression. Second, the reduced relative steady-state current demonstrates that altered gating also reduces current magnitude, likely through enhanced channel inactivation. Our data do not rule out the possibility that hERG1NP differentially regulates hERG1 subunits: hERG1a-1c and hERG1USO. This is particularly relevant given the upregulation of the hERG1NP in immature myocytes and the dynamic nature of hERG1 subunit abundance in the developing myocardium (4, 47, 48) and heart failure (23). hERG1 variants are enriched in cases of sudden death in the young (6, 7) and hERG1NP dysfunction could be a contributing factor in these cases.

Our data likely rule out a downstream “cryptic” promoter and/or IRES-mediated translation as drivers for hERG1NP expression. We generated the KCNH2-null stem cell line with a two base pair insertion in exon 6 (hERG1a numbering), which fully abolished hERG1NP expression. Any promoters downstream of the insertion would be predicted to remain functional. This suggests that proteolytic cleavage of the full-length channel generates the hERG1NP. And because the effects of the hERG1NP on channel gating were dependent upon nuclear targeting, these data suggest that the hERG1NP alters gene expression to indirectly modulate hERG1 channel function.

Our data identify a KCNH2-encoded peptide that does not, at least directly, modulate hERG1 function. This adds to a growing body of work that highlights roles for KCNH2 that are unrelated to cardiac repolarization. Recent work in rat cells demonstrated that rERG1 complexes with integrin β1 to activate FAK signaling during cardiac differentiation and cardiac recovery following sepsis (49, 50). hERG1 displays a similar role in cancer lines, where hERG1 promotes proliferation and metastasis (51, 52). Our data are interesting because hERG1NP expression diminished sharply with cardiomyocyte maturation, which roughly corresponds with the transition from hyperplasia (embryonic & fetal cardiomyocytes) to hypertrophy (neonatal & adult cardiomyocytes) (53). This suggests a potential role for the hERG1NP in cardiomyocyte proliferation. Indeed, the full-length C-terminus displayed a bimodal distribution (nuclear vs cytoplasmic targeting, c.f. Fig. 4). These HEK293 data also support cell-cycle as a factor in hERG1NP nuclear targeting.

hERG1’s widespread expression and association with diseases across multiple organs and tissues point to a fundamental role for KCNH2 in human physiology. hERG1 is widely expressed in the brain and its dysfunction is associated with schizophrenia (54, 55) and epilepsy (56, 57). Pancreatic beta cells express KCNH2, and loss-of-function KCNH2 variants are associated with hyperinsulinemia and diabetes (58, 59). KCNH2 is also expressed in smooth muscle tissues where it regulates contractile function (60, 61), and is associated with vascular remodeling (62, 63).

The integral role of ERG1 is best highlighted by work in mice. IKr minimally regulates adult murine cardiac excitability, yet loss of KCNH2 expression is embryonically lethal, triggering a myriad of morphological cardiac defects (64–66). KCNH2 transcripts are observed at the earliest stages of mouse cardiac development (67), and have even been reported in developing mouse embryos prior to implantation (68). In fact, loss-of-function KCNH2 variants have been identified in cases of tetralogy de Fallot (69, 70), which suggests that hERG1 may also contribute to cardiomorphogenesis in humans as well. Evolutionary analysis also demonstrates an ancestral origin for ERG channels that predates the divergence of cnidarians and bilaterians (71). With such an ancient origin it is rational to presume that hERG1 function is integrated in widespread aspects of physiology. The identification of hERG1NP represents one step closer to understanding the breadth of mechanisms by which KCNH2 regulates human physiology.

Finally, hERG1 is not the ion channel-related peptide to display a nuclear-targeted subdomain. Proteins encoded by GJA1 (Cx43), CACNA1A (CaV2.1), CACNA1C (CaV1.2), TRPM7 (TRPM7), and SCN1B (NaV channel β1 subunit) display similar activity (72–75). These peptides are generated by either proteolytic cleavage (72, 76–78) or alternative forms of translation such as a cryptic promoter or internal ribosomal entry site (73, 74, 79, 80). For example, mRNA of the CACNA1C gene encodes the full-length channel, CaV1.2, and a smaller, 75 kDa transcription factor called CCAT (73, 79). CCAT is identical to the C-terminus of CaV1.2 but is targeted to the nucleus where it regulates expression of neuronal signaling genes (73, 79). Similarly in CACNA1A, an internal ribosomal entry site (IRES) initiates translation of a transcription factor called α1ACT, which promotes neural and Purkinje cell development (74). The TRPM7 displays tissue-specific proteolytic cleavage. Similar to KCNH2, TRPM7 displays low level expression throughout the body, yet its disruption is embryonically lethal (72). The cleaved TRPM7 polypeptides phosphorylate core histones leading to altered gene expression that is necessary for embryonic development (81). Nuclear trafficking of these ion channel subdomains appears to function as an additional feedback mechanism between activity at the surface membrane and gene expression within the nucleus (73, 75, 79, 82). This growing body of research demonstrates that the roles of ion channel proteins limit aspect of cellular physiology beyond membrane potential.

## MATERIALS AND METHODS

### CRISPR KCNH2 deletion

KCNH2 deletion was completed by CRISPR by the University of Michigan Stem Cell and Gene Editing Core in the H9 human embryonic stem cell background. H9 cells were obtained from the WiCell stem cell repository. The gene-editing core generated a guide RNA (gRNA) targeting KCNH2-exon 6 using the CRIPSR design tool via the UCSC genome browser on the human GRCh38/hg38 assembly (https://genome.ucsc.edu/), sgRNA1: ATGAGGTCCACCACAGCCAGCGG, sgRNA2: CTCCTCGTTGGCATTGACGTAGG. We verified a two base pair deletion in exon 6 by Sanger sequencing using the forward and reverse primers TCCTCTCCCTACACCACCTG and CTCCTCCTCATTCTGCTTGG, respectively (Supplementary Fig. S1). Isogenic control and gene-edited KCNH2-null hESCs displayed appropriate pluripotency markers and differentiated into contracting hESC-CMs (Fig. 2 & Supplementary Fig. S1).

### Stem Cell Culture and Cardiac Differentiation

We cultured and differentiated human iPS and ES cells into cardiomyocytes using the GiWi protocol, as described (42). Briefly, we seeded stem cells on Matrigel-coated plasticware with iPS-brew medium. We checked media daily to remove spontaneous differentiation and passed the cells at 70% confluence. At the day of cell passage, we re-seeded cells to continue the line, or seeded the cells to grow monolayers for cardiac-directed differentiation. We plated 4×105 cells into each well of a 6-well plate and cultured them to ~80% confluence for treatment with GSK3 inhibitor for induction of mesodermal differentiation (day 0, D0). Following mesodermal differentiation, we treated cells with a Wnt inhibitor for induction of cardiac mesoderm (D2). On D4 we removed Wnt inhibition to direct the cells into cardiac progenitor cells. Cardiomyocytes with autonomous contractility emerged eight to ten days after initiation of cardiac-directed differentiation. We cultured the cardiomyocytes until 20 days after initiation of differentiation, and isolated purified cardiomyocytes by magnetic-beads assisted isolation with an iPSC-Derived Cardiomyocyte Isolation Kit, human (Miltenyi Biotec, USA) following the manufacturer’s recommendations. We plated the purified cardiomyocytes on Matrigel-coated coverslips for seven days before completing experiments.

### HEK293 Cell Culture

We maintained all cells at 37°C and 5% CO2 in a Heracell incubator (Thermo Fisher). We cultured HEK293 cells in minimum essential medium (MEM, Invitrogen, Cat. No. 11095080) supplemented with 10% fetal bovine serum (Thermo Fisher, Cat. No. SH30070.03) and split every 3-5 days at 60%-80% confluency.

### Stem Cell-Derived Cardiomyocyte Electrophysiology

We completed all recordings at physiological temperature (37 ± 1°C) using whole-cell patch clamp with an IPA Amplifier running and Sutterpatch (Sutter). We performed leak subtraction off-line based on measured current observed at potentials negative to IKr activation. The inter-pulse duration for all recordings was 10 seconds.

We sampled data at 10 kHz and low-pass filtered at 1 kHz. Cells were perfused with extracellular solution containing (in mM): 150 NaCl, 5.4 KCl, 1.8 CaCl2, 1 MgCl2, 15 glucose, 10 HEPES, 1 Na-pyruvate, and titrated to pH 7.4 using NaOH. Recording pipettes had resistances of 2-4.5 MW when backfilled with intracellular solution containing (in mM): 5 NaCl, 150 KCl, 2 CaCl2, 5 EGTA, 10 HEPES, 5 MgATP and titrated to pH 7.2 using KOH. We isolated IKr as an E-4031-sensitive current, as described (19, 35, 83). We preceded IKr recordings with a 100 ms step to −40 mV to inactivate the voltage-gated sodium currents. To elicit IKr we stepped from the −40 mV pre-pulse to membrane potentials between −50 mV and +60 mV in ten mV increments. We measured tail currents during a −40 mV, 3-second test pulse. To describe the voltage dependence of IKr activation, we normalized peak tail current to cellular capacitance, plotted current density as a function of pre-pulse potential, and fitted the data with the following Boltzmann equation:

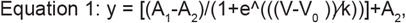

where A_1_ and A_2_ represent the maximum and minimums of the fit, respectively, V is the membrane potential, V_0_ is the midpoint, and k is the slope factor.

### HEK293 Electrophysiology

We completed all HEK293 recordings identical to cardiomyocyte recordings with the following exceptions. We completed HEK293 recordings at room temperature. We completed leak subtraction off-line at potentials −80 mV. To elicit I_hERG1_ we stepped from −80 mV to membrane potentials between −80 mV and +50 mV in ten mV increments. We measured tail currents during a −50 mV, 3-second test pulse. To describe the voltage dependence of IhERG1 activation, we normalized peak tail current to cellular capacitance, plotted current density as a function of pre-pulse potential, and fitted the data with a Boltzmann equation (Eq. 1). To measure the time course of deactivation we fit current decay during a −50 mV test pulse from +30 mV with a double exponential function:

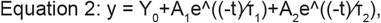

where Y_0_ is the asymptote, A_1_, and A_2_ represent the the relative components of the fast and slow time constats of decay (τ_1_ and τ_2_), respectively.

### Stem Cell-Derived and Native Cardiomyocyte Immunocytochemistry

We validated the differentiated cardiac lines using immunocy to chemistry targeting actin (phalloidin, cat #A12379 ThermoFisher) to display the cardiac sarcomere organization, and patch clamp electrophysiology measuring cardiac IKr, indicative of hERG1 expression. To target the hERG1 p-loop, we immunolabeled all cardiomyocytes with 1:200 dilution of primary antibody #ALX-804-652-R300 from Enzo Life Sciences and 1:250 dilution of secondary antibody goat anti-mouse Alexa Fluor 568 (#A-11004, Invitrogen). To target the hERG1 C-terminal domain we immunolabeled cardiomyocytes with a 1:150 dilution of primary antibody #ALX-215-049-R100 from Enzo Life Sciences and 1:250 dilution of secondary antibody AF647, Goat Anti-Rabbit, Cat. #4050-31 from Southern Biotech. We labeled the nuclei using 1:1000 dilution of DAPI (1μg/ml) for 15 minutes (ThermoScientific, Cat. #62248).

#### HEK293 Cell Immunocytochemistry

We cultured cells to 60% confluency in 6-well plastic dishes. We transfected cells with 1 μg DNA using Lipofectamine 3000. 24 hours after transfection, we replated cells onto 12 mm No. 1.5 glass coverslips. 48 hrs after transfection, we fixed cells using 4% pfa for 15 minutes. We washed the fixed cells three times using PBS and stained using DAPI at (1μg/ml) for 15 minutes and then washed three more times with PBS. We mounted the coverslips on microscope slides using Prolong Gold mounting medium and completed imaging 48 hours after mounting. We completed all imaging using a Zeiss 880 confocal microscope.

#### DNA constructs

The hERG1 proximal domain (666-872-Citrine. pcDNA3), distal domain (873-1159-Citrine.pcDNA3), and full-length C-terminal domain (666-1159-Citrine.pcDNA3) plasmids were generously provided by Professor Matthew Trudeau at the University of Maryland Medical School. We generated the 873-1159-Citrine ΔNLS mutant using site-directed mutagenesis (QuikChange II Kit, Agilent Technologies) using the distal plasmid as a template. We designed the primers (Forward, 5’-ccgtgcgcctactgaagccaccctctaac-3’: Reverse, 5’-gttagagggtggcttcagtaggcgcacgg-3’) with the QuikChange Primer Design (agilent.com) and the primers were synthesized by Eurofins. We verified the sequences of all constructs by DNA sequencing (Eurofins).

### Statistical Analysis

We completed analysis using IgorPro and Prism (Graphpad). We evaluated values for normality and outliers before statistical evaluation. All data are reported as mean ± SEM and we compared values using an ANOVA and Bonferroni post hoc t-tests. Statistical significance was taken at p < 0.05.

## ACKNOWLEDGMENTS

This work was supported by R00HL133482 and a Samuel and Jean Frankel Cardiovascular Center Amplifier Grant to DKJ, T32-GM140223 to FGSC, and T32-HL125242 to CU. The authors thank the University of Michigan Gene Editing Core for generating the KCNH2-null human embryonic stem cell line, and Matt C. Trudeau for providing the hERG1 C-terminal constructs.

## Notes

### Competing Interest Statement

The authors have declared no competing interest.

