## Supplementary Figure S1 for "*KCNH2* encodes a nuclear-targeted polypeptide that mediates hERG1 channel gating and expression"

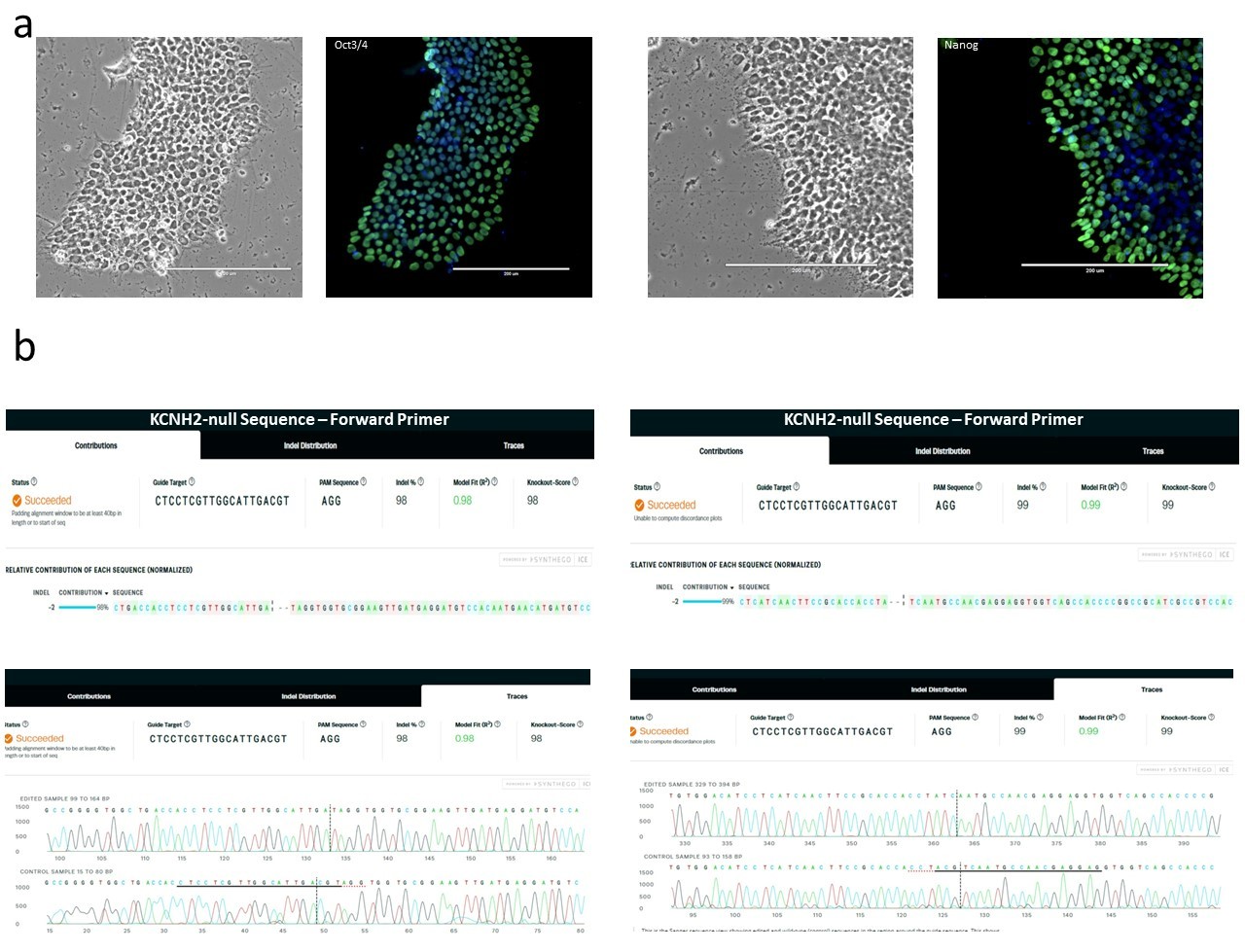

**Supplementary Figure S1.** Pluripotency markers and gene sequence of CRISPR-edited H9 human embryonic stem cells. (a) Brightfield and immunofluorescent images depicting *KCNH2*-null human embryonic stem cell colonies immunostained for the pluripotency markers Oct3/4 (*left*) and Nanog (*right*). (b) Forward (*top left*) and reverse (*top right*) primers used to sequence the target site within exon 6 of *KCNH2*. The corresponding gene sequence from each primer, depicting the 2 bp insertion, is shown below.
